# The yeast core spliceosome maintains genome integrity through R-loop prevention and α-tubulin expression

**DOI:** 10.1101/272807

**Authors:** Annie S. Tam, Veena Mathew, Tianna S. Sihota, Anni Zhang, Peter C. Stirling

## Abstract

To achieve genome stability cells must coordinate the action of various DNA transactions including DNA replication, repair, transcription and chromosome segregation. How transcription and RNA processing enable genome stability is only partly understood. Two predominant models have emerged: one involving changes in gene expression that perturb other genome maintenance factors, and another in which genotoxic DNA:RNA hybrids, called R-loops, impair DNA replication. Here we characterize genome instability phenotypes in a panel yeast splicing factor mutants and find that mitotic defects, and in some cases R-loop accumulation, are causes of genome instability. Genome instability in splicing mutants is exacerbated by loss of the spindle-assembly checkpoint protein Mad1. Moreover, removal of the intron from the α-tubulin gene *TUB1* restores genome integrity. Thus, while R-loops contribute in some settings, defects in yeast splicing predominantly lead to genome instability through effects on gene expression.

## INTRODUCTION

Maintenance of a stable genome is a complex process involving the coordination of essentially all DNA transactions, including transcription, chromatin state, DNA replication, DNA repair and mitosis. Regulation of genome stability is critical to prevent cancer (Hanahan and Weinberg, 2011). While screens across model organisms and human cells have implicated numerous genes as regulators of genome maintenance, in many cases we do not understand the mechanism of action (Paulsen et al., 2009; Stirling et al., 2011).

Defects in RNA processing have been implicated in genome instability across species, and in both human cancer and repeat-expansion diseases (Richard and Manley, 2017). Indeed, RNA splicing factors are frequently lost or mutated in various cancers where they shift gene expression landscapes, favoring oncogenesis (Darman et al., 2015; Dolatshad et al., 2015; Joshi et al., 2017). The detailed role of splicing in preventing genome instability is still unclear. Previous work has suggested that loss of splicing factors like SF2/ASF (Li and Manley, 2005), or treatment with splicing inhibitors (Wan et al., 2015), cause the accumulation of DNA:RNA hybrids in genomic DNA. These three-stranded R-loop structures are known to contribute to genome instability by exposing single-stranded DNA (ssDNA) and by blocking replication fork progression, causing replication stress induced genome instability (Aguilera and Garcia-Muse, 2012; Chan et al., 2014b). Still other groups have suggested that splicing factor disruption, such as loss of CDK12, causes changes in gene expression, which reduce the activity of canonical genome maintenance factors leading to defects in DNA repair (Blazek et al., 2011). The mechanisms that might predominate when splicing is disrupted are not clear.

We previously identified a large number of RNA processing factors whose disruption in yeast leads to genome instability, including core components of all small nuclear ribonucleoprotein (snRNP) complexes involved in splicing (Stirling et al., 2011). Importantly, only 5% of yeast genes encode introns, which reduces the complexity of interpreting specific splicing changes as drivers of genome instability (Parenteau et al., 2008). Here we set out to resolve the contribution of R-loops versus gene expression changes in splicing-loss induced genome instability by directly testing the mechanisms of genome instability in mutants disrupted for the U1, U2 and U4/U6.5 snRNPs. While we observe evidence of R-loop induced DNA damage in some splicing mutants, each splicing mutant appears to cause aberrant splicing of the α-tubulin transcript from the *TUB1* gene. Ensuing mitotic defects arising from Tub1 depletion are therefore a common driver of chromosome loss in yeast splicing mutants. Thus, informational defects in the transcriptome arising from splicing defects are a major route to genome instability.

## RESULTS and DISCUSSION

### U1, U2 and U4/U6.U5 splicing factor mutations lead to chromosome instability

Previous screens have identified at least 25 splicing proteins that, when disrupted in yeast, lead to chromosomal instability (CIN) (Stirling et al., 2011). To begin to understand whether R-loops or other mechanisms drove genome instability, we conducted CIN assays in strains with mutations in each of the core snRNP complexes involved in establishing the splicing reaction (Measday and Stirling, 2015). Since each spliceosomal snRNP is essential we used temperature-sensitive (ts) alleles of *YHC1* (U1), *HSH155* (U2), and *SNU114* (U4/U6.U5), each of which had strong splicing defects as measured with a LacZ splicing reporter (Galy et al., 2004) or at an endogenous spliced transcript (**Figure S1A** and **S1B**). In all three mutants, we observed increases in artificial chromosome loss by the chromosome transmission fidelity (ctf) assay (Measday and Stirling, 2015) and by plasmid loss assays (**Figure 1A** and **1B**). To test the stability of endogenous chromosomes we monitored chromosome III copy number with an integrated LacO-array in cells expressing LacI-GFP to create a green dot marked chromosome III. In unbudded G1 cells all three splicing mutants showed an increased rate of gain of a LacI-GFP marked chromosome III, suggesting that a chromosome gain event has taken place (**Figure 1C**). Thus, splicing mutants led to a significant increase in chromosomal instability across different assays from plasmids to native chromosomes.

**Figure 1.**
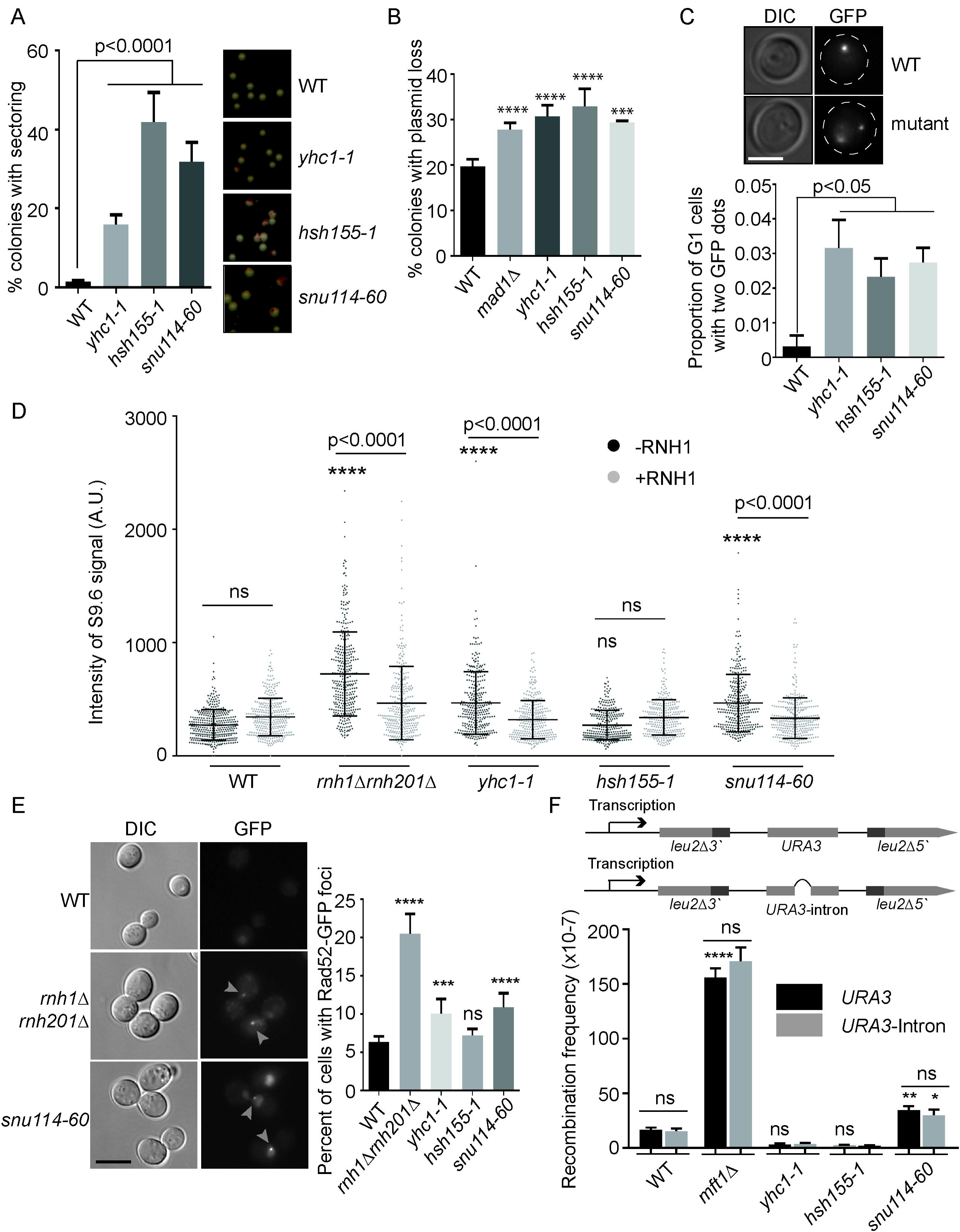
Genome instability phenotypes of splicing mutants. (A) CTF phenotypes in *yhc1-1* (U1), *hsh155-1* (U2), and *snu114-60* (U4/U6.U5) alleles are indicated by the percent of red sectored colonies. Representative images of each strain displayed on right. (B) Plasmid loss frequency in splicing alleles. Deletion of spindle assembly checkpoint gene *MAD1* was used as a positive control. (C) Frequency of unbudded cells with >1 GFP-marked LacO array indicating gain of chromosome III. Top: representative images showing one or two GFP-marked LacO array in WT or mutant, respectively. Dashed lines indicate cell outlines (D) s9.6 antibody DNA:RNA hybrid staining intensities of splicing mutant chromosome spreads. Asterisks indicate p-values relative to WT-RNH1. (E) Frequency of Rad52-GFP DNA damage foci in the indicated strains. Damage foci are highlighted in GFP channel (grey arrows). (F) Frequency of direct repeat recombination on the indicated plasmid. A schematic of each reporter construct is presented above panel. Deletion of THO complex subunit *MFT1* was used as a positive control. Asterisk above *snu114-60 URA3*-intron is the p-value relative to WT *URA3*-intron. For all, mean values with S.E.M. error bars are shown, *n = 3*. (A-C, E) Fisher’s exact test (F) Student’s t-test (D) one-way ANOVA. *p<0.05; **p<0.01; ***p<0.0005; ****p<0.0001. Scale bar in C = 2μm; scale bar in E =5μm.

### Sporadic R-loop accumulation and DNA damage phenotypes in splicing mutants

The CIN phenotypes observed in splicing mutants could arise by several mechanisms including the formation of transcription coupled R-loops. To test this model we first performed chromosome spreads and used the S9.6 antibody to detect DNA:RNA hybrids using immunofluorescence (Wahba et al., 2011). Interestingly, while *yhc1-1* and *snu114-60* alleles showed high levels of R-loop accumulation, the *hsh155-1* allele had no increase in R-loops (**Figure 1D**). This suggests that R-loops cannot be a common mechanism of instability across mutants. Indeed, when we analyzed Rad52-GFP foci, a marker of DNA damage repair that should increase if R-loops are driving genome instability, we found that only *yhc1-1* and *snu114-60* had a significant increase in Rad52 foci (**Figure 1E**). To further test phenotypes known to correlate with aberrant R-loop levels, we used a plasmid-based direct repeat recombination system to test for hyper-recombination and only *snu114-60* showed a significant increase in recombination using both plasmid and integrated genomic reporters (**Figure 1F and Figure S1D**). Importantly, both DNA damage and recombination phenotypes in *snu114-60* were suppressed by ectopic expression of RNaseH1, confirming that R-loops can indeed play a role in genome instability for some splicing mutants (**Figure S1C and Figure S1D**). These data confirm that R-loops are not a unifying mechanism of genome instability across the various spliceosomal snRNP mutants. Indeed, R-loop accumulation was only seen in a subset of splicing mutants in a previous screen, suggesting splicing mutant allele specificity that we currently do not understand (Chan et al., 2014a).

### Genetic interaction profiling reveals mitotic defects in hsh155-1

Mutations in Hsh155 exhibited strong CIN phenotypes, but showed no evidence of increased DNA damage or R-loops. We hypothesized that if a common mechanism of CIN existed for splicing mutants, it would be at play in *hsh155-1* alleles. To determine this function, we performed a synthetic genetic array (SGA) screen using *hsh155-1* as a query strain. This screen identified 102 negative and 103 positive genetic interaction candidates (**Table S1**). Interacting genes grouped according to their function identified the expected negative interactions with other RNA splicing and transcription associated genes. Indeed, a greater than expected number of essential intron-containing genes were negative interactors, consistent with a splicing defect enhancing phenotypes of these mutant genes (**Table S2**). Positive interactions with proteasome subunits or translational apparatus may reflect stabilization of mutant Hsh155 protein leading to healthier cells (**Table S1**) (van Leeuwen et al., 2016). A clear connection to chromosome segregation came when analyzing gene ontology (GO) terms for the negative genetic interactions of *hsh155-1* (**Table S3**). GO term enrichments visualized with REVIGO (Supek et al., 2011) highlight the enrichment of processes like microtubule polymerization, and mitotic sister chromatid biorientation and components like the Mis12/MIND complex of the kinetochore (**Figure 2A** and **Figure 2B**). We validated by spot dilutions that *hsh155-1* have negative interactions with mitotic genes like cohesin (*MCD1*), core kinetochore subunits (*MIF2*) and spindle regulators (*STU1*) (**Figure 2C**). We further used growth curves to identify subtle changes in growth in the *hsh155-1* double mutants (**Figure 2D**). Analysis of published SGA profiles for *yhc1-1* and *snu114-60* (www.thecellmap.org) also revealed links to the mitotic apparatus (Costanzo et al., 2016), which encouraged us to explore mitotic defects in all three alleles.

**Figure 2.**
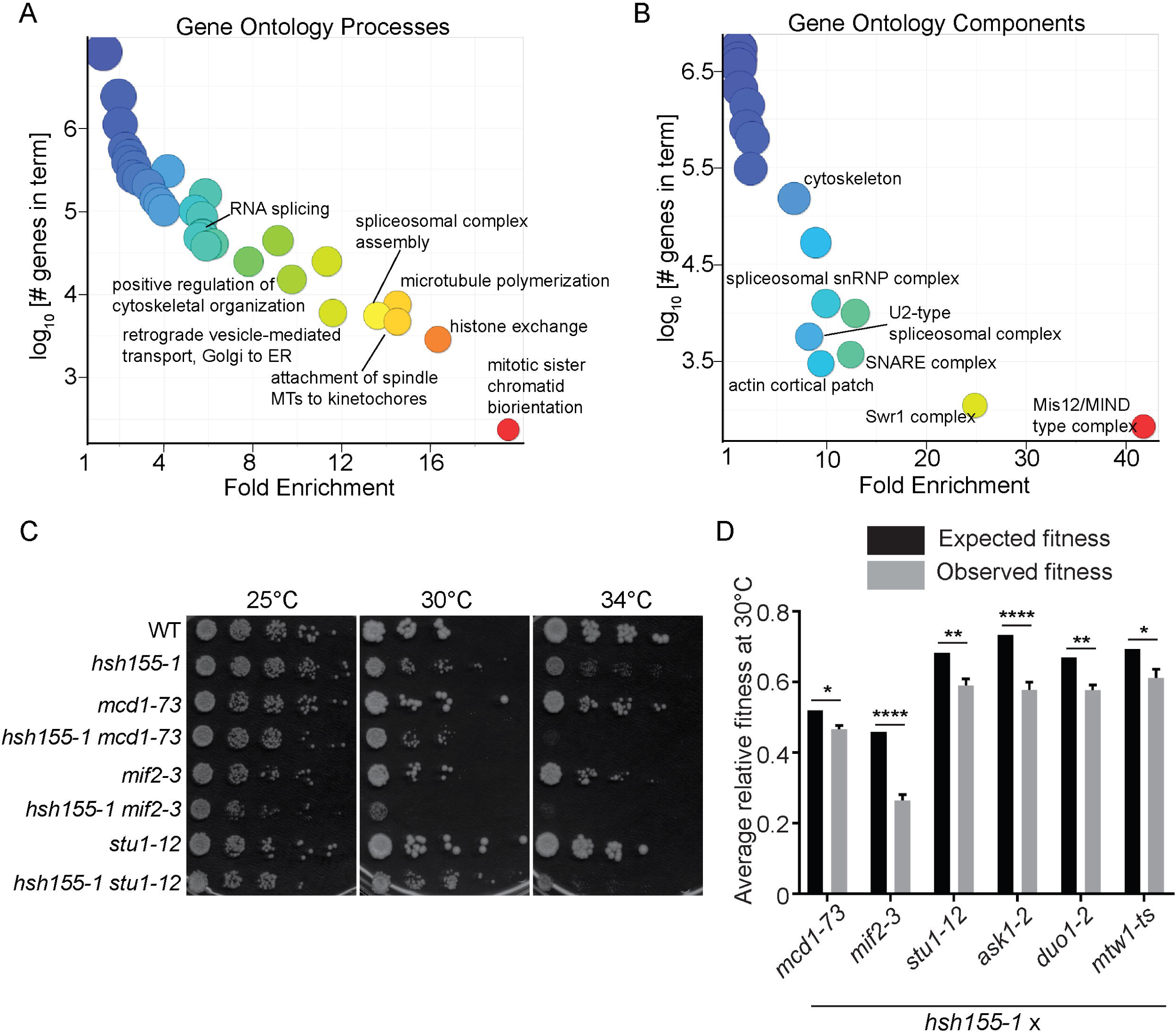
Genetic interaction network of *hsh155-1*. (A) GO biological process and (B) cellular component enrichments for *hsh155-1* negative interactions. The complete list is in **Table S2**. REVIGO was used to visualize a reduced set of terms by the number of genes associated with the term, and the fold enrichment. Warmer colors associate with higher enrichment. Validation of negative genetic interactions with kinetochore, cohesin and spindle mutants by (C) spot dilution assays at the indicated temperatures and (D) growth curves at 30°C, graphical results of which indicate observed and expected fitness values.*p<0.05 **p<0.01 ****p<0.0001.

### Spliceosome mutants have mitotic defects

To test potential mitotic defects in our splicing mutants, we first measured the cell cycle distribution of cells by budding index, using Hta2-mCherry as a nuclear marker. After a shift to a non-permissive temperature of 37°C, a significant proportion of the splicing mutants accumulated as large-budded G2/M cells, indicating a potential mitotic delay and defect (**Figure 3A**). These results complemented our SGA screen (**Figure 2**) highlighting negative genetic interactions between *hsh155-1* and genes with functions in spindle and kinetochore subunits. To further test how the spindle could be influencing genetic instability, we deleted the spindle assembly checkpoint (SAC) component *MAD1* in each splicing mutant. Loss of *MAD1* further sensitized *hsh155-1* and *snu114-60* alleles to the microtubule depolymerizing drug benomyl at semi-permissive temperatures (**Figure 3B**). We did not detect enhanced benomyl sensitivity in *yhc1-1* using this assay, possibly because of limited allelic penetrance at 30°C. By quantifying the rate of endogenous chromosome III loss using the A-like faker assay (Novoa et al., 2018) in the single and double mutants at permissive temperature of 25°C, we found a dramatic synergy between disruption of *SNU114* or *HSH155* and loss of the SAC (**Figure 3C**). Recurrently, *yhc1-1* phenotype was not enhanced, consistent with the lack of benomyl sensitivity under the same conditions and likely a limitation of the ts-allele. Overall, these data are consistent with a mitotic defect in splicing mutants that additionally requires the activity of the SAC for genome maintenance, in particular for the alleles tested in the U2 (*hsh155-1*) and tri-snRNP (*snu114-60*) complexes.

**Figure 3.**
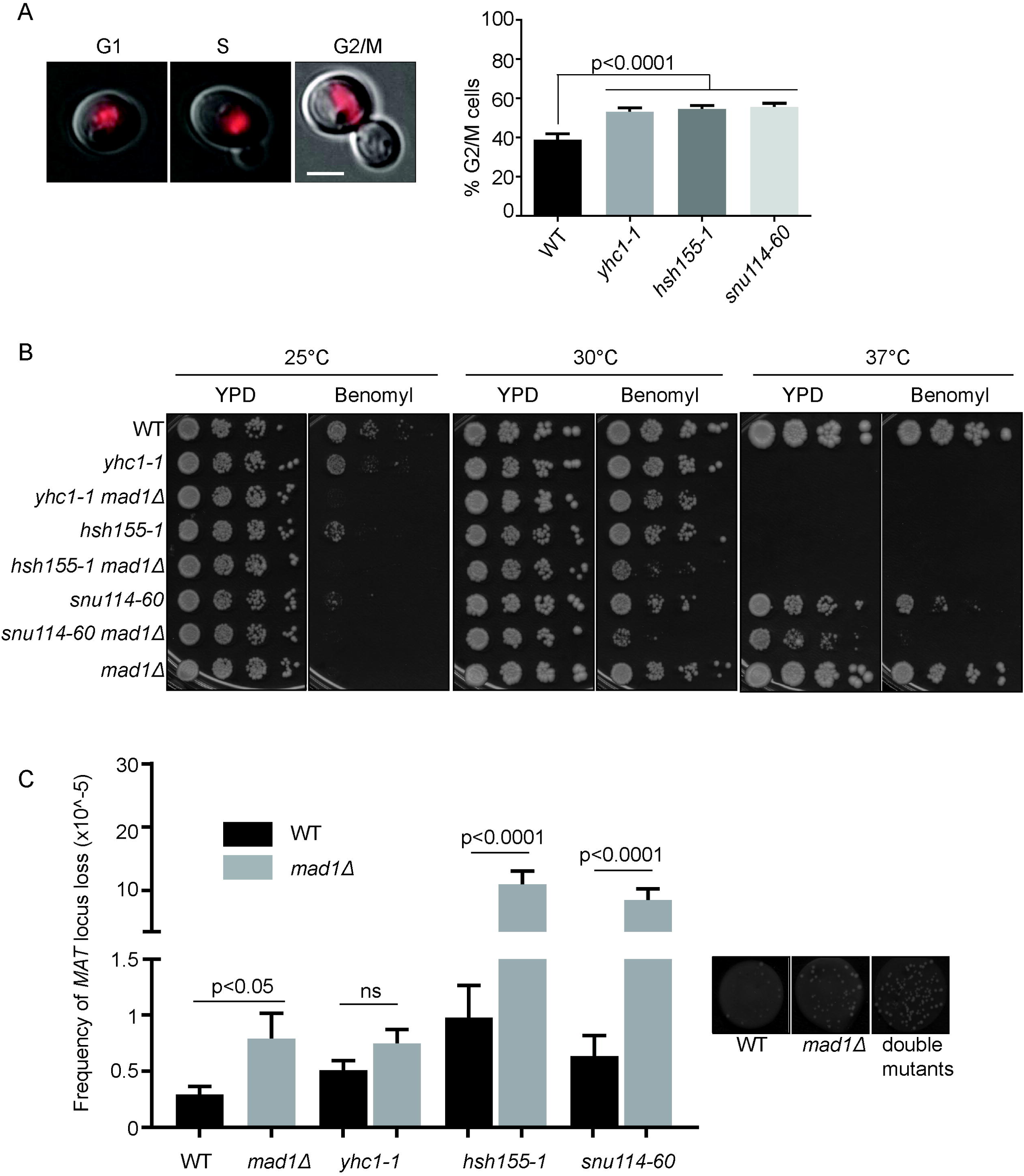
Mitotic defects in spliceosome mutants. (A) Proportion of G2/M cells visualized under the microscope. Position of nuclei was determined by Hta2-mCherry fluorescence (left panels, scale bar = 2μm). Fisher’s exact test; Mean values with S.E.M. error bars are shown, *n = 3*. (B) Benomyl sensitivity, relative to DMSO control determined by spot dilution assay for the indicated mutants and temperatures. (C) Frequency of *MAT* locus loss in single (black bars) and *mad1Δ* double mutants (grey bars), grown at 25°C. Representative images of WT, *mad1Δ* and double mutant colonies shown on right. Student’s t-test; Mean values with S.E.M. error bars are shown, *n = 3*.

### Tubulin levels control genome integrity in splicing mutants

Only ∼5% of yeast genes are spliced and the bulk of splicing flux is accounted for by the production of ribosomal proteins, many of which are encoded by spliced genes (Parenteau et al., 2008). The cytoskeleton is also enriched for intron-containing genes. Indeed, splicing of the *TUB1* transcript, encoding α-tubulin, has previously been implicated in cell cycle delays in other splicing mutants, although never directly in genome stability (Burns et al., 2002; Dahan and Kupiec, 2002). The effects on Tub1 protein was tested by western blot and a decrease in Tub1 protein levels was observed for all three splicing mutants as temperature increased (**Figure 4A**). The quantitative PCR of *TUB1* mRNA transcript levels under the same conditions indicated a mild decrease in *TUB1* mRNA expression in *hsh155-1* and *snu114-60.* Interestingly a dramatic increase in intron retention was observed for all three mutants (**Figure 4B**), supporting the notion that defective *TUB1* mRNA splicing drives loss of protein expression. To directly connect defective splicing-induced aberrant α-tubulin levels to genome maintenance, we retested CIN in splicing mutants encoding an intronless *TUB1* gene. As expected, intronless *TUB1* (*tub1*Δ*i*) increased the amount of Tub1 protein expressed in each splicing mutant, as measured by western blot (**Figure 4A**). More importantly, intronless *TUB1* partially suppressed the CTF phenotype observed in each splicing mutant, although the effects in *yhc1-1* were mild (**Figure 4C**). Since the CTF assay measures loss of an extrachromosomal fragment, we also tested the effect of intronless *TUB1* on the rate of endogenous chromosome III mis-segregation using the LacO-LacI-GFP system and found clear suppression of aneuploidy in all three splicing mutants (**Figure 4D**). Thus, when splicing of *TUB1* fails, Tub1 protein is reduced, stressing the mitotic apparatus and reducing mitotic fidelity in a way that is buffered by the SAC (**Figure 4E**).

**Figure 4.**
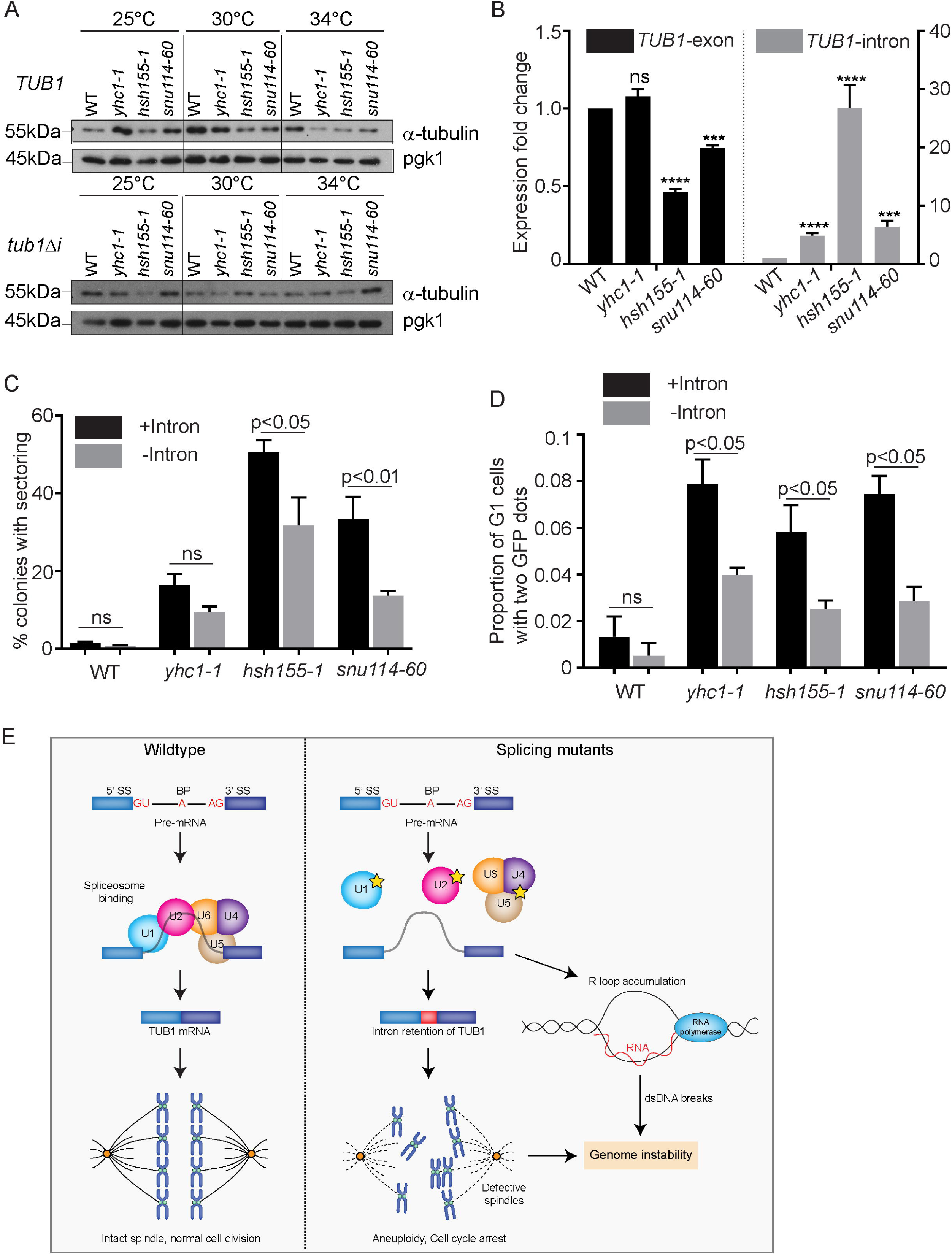
Tubulin stability contributes to genome maintenance in splicing mutants. (A) Western blot indicating α-tubulin protein levels in splicing mutants at increasing temperature. Top: *TUB1*; Bottom: intronless *TUB1* (B) Quantification of *TUB1* mRNA transcript levels from exon region in WT, *yhc1-1, hsh155-ts* and *snu114-60* (left) and *TUB1* intron region (right) by reverse transcription-quantitative PCR normalized to *SPT15* and relative to WT. Mean values with S.E.M. error bars are shown, *n = 3*. Asterisks show p-values of ΔΔCt – ***p=0.0002; ****p<0.0001. (C) CTF phenotypes in *hsh155-1* and *snu114-60* alleles partially suppressed with intronless *TUB1* (grey bars). (D) Endogenous chromosome stability partially restored in *yhc1-1, hsh155-1* and *snu114-60* with intronless *TUB1* (grey bars). (C,D) Fisher’s exact test; Mean values with S.E.M. error bars are shown, *n = 3*.(E) Model of defective splicing-induced genome instability.

### Perspective

Determining sources of genome instability is important to understand the accumulation of mutations during adaptation and in human disease. While much is known about direct genome maintenance factors in DNA replication, repair and mitosis, considerably less is known about the effects of other cellular pathways. Nonetheless, these non-canonical pathways account for a large proportion of reported genome maintenance factors (Stirling et al., 2011). RNA processing has emerged as a major contributor to genome maintenance and various mechanisms have been described. Direct roles for some RNA processing factors have been found in DNA repair, such as the moonlighting function of Prp19 in ATR activation (Marechal et al., 2014), or the role of the spliceosome in R-loop mediated ATM activation (Tresini et al., 2015). More recently, dominant cancer-associated splicing mutations have been linked to RNA polymerase pausing and R-loop accumulation (Chen et al., 2018). Still other studies have suggested a role for the aberrant gene expression landscapes produced in RNA processing mutants as drivers of genome instability (Blazek et al., 2011; Vohhodina et al., 2017). Here we test representative core snRNP mutants of the spliceosome that were previously implicated in CIN. Our findings indicate that R-loops are not universally formed as a common source of instability in splicing mutants. Instead, impaired splicing and thus expression of a key microtubule protein, Tub1, is a common driver of chromosome loss phenotypes in yeast splicing mutants. Importantly, restoration of Tub1 by removing the intron partially rescued the CIN phenotype of the splicing mutants. The partial rescue suggests that other mechanisms are also contributing to genome instability. Indeed for *snu114-60*, we find that both R-loop accumulation and changes in Tub1 protein levels clearly contribute to genome instability. What regulates the penetrance of a specific splicing mutant for either R-loop or gene expression based CIN phenotypes is likely to be allele specific, and so far remains unclear. Broadly speaking this study suggests that as splicing mutants are now frequently seen in various cancers, interpretation of their potential effects on genome maintenance needs to account for both potential direct R-loop mediated damage, and indirect changes in the translatome. If, and how, aberrant R-loop accumulation and gene expression changes are connected in cells with defective splicing is an important area for future study.

## EXPERIMENTAL PROCEDURES

### Yeast strains, growth and CIN assays

All yeast strains were in the s288c background (**Table S4**), and were grown under standard conditions in the indicated media and growth temperature. For benomyl sensitivity, strains were compared between YPD+2% DMSO (control) or YPD+15μg/mL benomyl (Sigma-Aldrich cat#45339). Growth curves were conducted in a Tecan M200 plate reader and compared using the area under the curve as previously described (Chang et al., 2017). The CTF and ALF assays were performed as described (Novoa et al., 2018; Stirling et al., 2011). For plasmid loss, strains carrying *pRS313::HIS3* were grown overnight in SC-histidine, then plated on YPD and allowed to form colonies without selection before replica plating onto SC-histidine. The frequency of colonies that could not grow on SC-histidine is reported as the plasmid loss rate. For all experiments significance of the differences was determined using Prism7 (GraphPad Software). For all experiments, sample means were compared with Fisher’s exact test, Student’s t tests or ANOVA for multiple comparisons as indicated.

### Synthetic Genetic Array and validation

SGA screening was performed as described previously for *URA3*-marked query strains (Costanzo et al., 2016; Stirling et al., 2011) and scored using the Balony software package (Young and Loewen, 2013). From raw colony scores, we chose a cut-off of p<0.05 and an experiment-control score of ≥|0.5| for list analysis and GO term enrichment using the Princeton generic GO term-finder (http://go.princeton.edu/cgi-bin/GOTermFinder) and visualized with REVIGO (Supek et al., 2011). *snu114-60* and *yhc1-1* interactions are available at http://thecellmap.org (Usaj et al., 2017). For hit validation, fresh double mutant strains were made by tetrad dissection and tested in quantitative growth curves, or by spot dilution assays. Observed area under the curve for double mutants was compared to a multiplicative model of the predicted fitness based on the fitness of the two single mutants.

### Image and cell cycle analysis

Imaging of budding index by differential interference contrast (DIC), or fluorescence of Rad52-GFP and GFP labeled chromosome III was conducted on a Leica DMi8 microscope using an HCX plan apochromat 1.4 NA oil immersion 100x lens. The images were captured at room temperature by an ORCA Flash 4.0 V2 camera (Hamamatsu Photonics), using MetaMorph Premier acquisition software (Molecular Devices). Scoring was done in ImageJ (National Institutes of Health). For Rad52 foci, all cells were scored as negative or positive for a focus. Imaging of LacO-LacI-GFP labeled chromosome III was performed as described (Woodruff et al., 2009). For scoring, first unbudded cells were selected in the DIC channel, then scored for the presence of 1 or ≥2 LacI-GFP dots. Chromosome spreads were performed exactly as described; primary DNA-RNA Hybrid [S9.6] (Kerafast cat#ENH001); secondary Alexa Fluor® 568 goat anti-Mouse IgG (Invitrogen cat#A-11004) (Wahba et al., 2011).

### Splicing efficiency assay

Splicing assay protocol was performed as described (Galy et al., 2004). All measurements were taken with individual transformants in triplicate. Cells were struck as a patch on SC-leucine and then replica plated to glycerol-lactate–containing SC medium without leucine (GGL-leu). Cells from each patch were inoculated in liquid GGL-leu media for 2 h at 30°C and then were induced with final 2% galactose for 4 h. Cells carrying reporters were lysed and assayed for β-galactosidase assay using a Gal-Screen β-galactosidase reporter gene assay system for yeast or mammalian cells (Applied Biosystems) as per the manufacturer’s instructions and read with a SpectraMax i3 (Molecular Devices). Relative light units were normalized to cell concentration as estimated by measuring OD600.

### Recombination assays

To construct the recombination plasmids, pRS314GLB (de la Loza et al., 2009) was linearized with *Bgl*II, and ligated with *URA3* and *URA3*-intron sequences with *Bgl*II sites (Aksenova et al., 2013). Following transformation and plasmid isolation, presence of *URA3* and *URA3*-intron was confirmed by sequencing. Direct repeat recombination assays were performed as described (Chang et al., 2017).

### Western blot

Whole-cell extracts were prepared by 100% trichloroacetic acid extraction (2×10^6^ cells). Lysates were separated on a 10% SDS-PAGE gel, transferred to nitrocellulose and probed with anti-Tub1 (Invitrogen cat#322500) (1:3000 dilution), or anti-Pgk1 (Abcam cat#38007) (1:500 dilution) as a loading control.

### RNA isolation, cDNA preparation, and reverse transcription–quantitative PCR analysis

Total RNA was isolated from 0.5-1 OD cell cultures shifted to 34°C for 3.5 h, using the yeast RiboPure RNA Purification kit (Ambion). 1 μg of cDNA was reverse transcribed using anchored-oligo(dT)18 primer and Transcriptor Reverse transcription (Roche). Reverse transcription–quantitative PCRs were performed and analyzed using SYBR green PCR Master Mix and a StepOnePlus Real-Time PCR system (Applied Biosystems).

## ACKNOWLEDGEMENTS

We thank Andres Aguilera, Hannah Klein, David Drubin, Benoit Palancade, Sherif Abou Elela, Douglas Koshland, and Philip Hieter for strains and plasmids. A.S.T. is a supported by the Cordula and Gunter Paetzold Fellowship. P.C.S. is a Canadian Institutes of Health Research (CIHR) New Investigator and Michael Smith Foundation for Health Research scholar. This work was supported by CIHR (MOP-136982) and the Natural Sciences and Engineering Research Council of Canada (RGPIN 2014-04490).

## SUPPLEMENTARY INFORMATION

**Figure S1**. Related to Figure 1. (A) Splicing efficiency of a LacZ reporter, values are relative to WT splicing level. Schematic of reporter constructs are presented above panel. Student’s t-test; Mean values with S.E.M. error bars are shown, n = 3. (B) Quantification of *RPL33B* mRNA transcript levels from intron region in WT, *yhc1-1, hsh155-ts* and *snu114-60* by reverse transcription-quantitative PCR normalized to *SPT15* and relative to WT. Mean values with S.E.M. error bars are shown, *n = 3*. P-value calculated from ΔΔCt levels (ANOVA). (C) Suppression of *snu114-60* Rad52-GFP DNA damage foci by ectopic *RNH1* expression (grey bars) compared to empty vector (black bars). ANOVA; Mean values with S.E.M. error bars are shown, *n = 3*. (D) Suppression of recombination frequency by ectopic *RNH1* expression (grey bars) compared to empty vector (black bars). ANOVA; Mean values with S.E.M. error bars are shown, *n = 3*. Schematic of reporter construct is presented above panel.

**Table S1**. Related to Figure 2. Raw SGA data for *hsh155-1*.

**Table S2**. Related to Figure 2. List of intron-containing genes identified as negative interactors in *hsh155-1* SGA screen.

**Table S3**. Gene Ontology terms enriched among *hsh155-1* negative interacting partners.

**Table S4**. Yeast strains and plasmids.

